# An insula-central amygdala circuit for behavioral inhibition

**DOI:** 10.1101/156216

**Authors:** Hillary Schiff, Anna Lien Bouhuis, Kai Yu, Mario A. Penzo, Haohong Li, Miao He, Bo Li

## Abstract

Predicting which substances are suitable for consumption during foraging is critical for animals to survive. While food-seeking behavior is extensively studied, the neural circuit mechanisms underlying avoidance of potentially poisonous substances remain poorly understood. Here we examined the role of the insular cortex (IC) to central amygdala (CeA) circuit in the establishment of such avoidance behavior. Using anatomic tracing approaches combined with optogenetics-assisted circuit mapping, we found that the gustatory region of the IC sends direct excitatory projections to the lateral division of the CeA (CeL), making monosynaptic excitatory connections with distinct populations of CeL neurons. Specific inhibition of neurotransmitter release from the CeL-projecting IC neurons prevented mice from acquiring the “no-go” response, while leaving the “go” response largely unaffected in a tastant (sucrose/quinine)-reinforced “go/no-go” task. Furthermore, selective activation of the IC-CeL pathway with optogenetics drove unconditioned lick suppression in thirsty animals, induced aversive responses, and was sufficient to instruct conditioned action suppression in response to a cue predicting the optogenetic activation. These results indicate that activity in the IC-CeL circuit is necessary for establishing anticipatory avoidance responses to an aversive tastant, and is also sufficient to drive learning of such anticipatory avoidance. This function of the IC-CeL circuit is likely important for guiding avoidance of substances with unpleasant tastes during foraging in order to minimize the chance of being poisoned.

**Significance Statement:** The ability to predict which substances are suitable for consumption is critical for survival. Here we found that activity in the insular cortex (IC) to central amygdala (CeA) circuit is necessary for establishing avoidance responses to an unpleasant tastant, and is also sufficient to drive learning of such avoidance responses. These results suggest that the IC-CeA circuit is critical for behavioral inhibition in anticipation of potentially poisonous substances during foraging.

## Introduction

The ability to suppress actions that can lead to harmful consequences is critical for survival. For example, animals, including humans, stop consummatory behavior when encountering food or liquid with an unpleasant taste, which indicates the existence of a potentially poisonous substance. Animals are also capable of learning to use environmental cues (such as an odor, color, location or context) to predict unpleasant properties (such as a bitter taste, toxicity) of a substance, and subsequently using these predictive cues to guide avoidance of the substance during foraging. While food approaching and reward seeking behaviors are extensively studied (Balleine, 2005, 2011; Kelley, 2004), the neural circuit mechanisms that underlie innate and learned suppression of actions that may lead to food poisoning, or aversive consequences in general, are poorly understood.

The insular cortex (IC), including the gustatory cortex (GC), plays an important role in processing taste and visceral information (Accolla and Carleton, 2008; Katz et al., 2001; Samuelsen and Fontanini, 2017; Yamamoto et al., 1985). The IC is also engaged in various learning tasks, especially when learning to associate a neutral tastant (also known as a conditioned stimulus, CS) with an intrinsically aversive stimulus (also known as an unconditioned stimulus, US), or to associate a non-gustatory CS with an appetitive or aversive tastant US (Bermudez-Rattoni, 2004; Kusumoto-Yoshida et al., 2015; Vincis and Fontanini, 2016; Yasoshima and Yamamoto, 1998). In one such task, a “go/no-go” task in which auditory “go” and “no-go” CSs predict the delivery of sucrose and quinine, respectively, specific CS-evoked responses develop in the IC as rats learn the predictive value of each CS (Gardner and Fontanini, 2014). IC neurons also show CS-evoked responses in a conditioned food-approaching task (Kusumoto-Yoshida et al., 2015). These findings indicate that associative learning driven by gustatory reinforcement induces plastic changes in the IC, and suggest that the IC may participate in guiding approach or avoidance responses during feeding or foraging behaviors. Consistently, pharmacological and optogenetic inhibition of the IC impairs conditioned food approaching behavior (Kusumoto-Yoshida et al., 2015). Nevertheless, whether the IC is also essential for avoidance of aversive tastes remains unknown.

The central nucleus of the amygdala (CeA), including its lateral division (CeL), is a prominent downstream structure of the IC. As it is also a direct recipient of taste information from the brainstem (Carter et al., 2013), the CeA is anatomically positioned to process convergent taste information. Indeed, it has been shown that the CeA encodes taste identification and palatability, responses that follow those in the IC (Sadacca et al., 2012). These findings suggest that the CeA may process and use the information originating from the IC to influence taste-motivated behaviors.

The CeL plays an important role in the learning and expression of defensive behaviors (Ciocchi et al., 2010; Goosens and Maren, 2003; Li et al., 2013; Wilensky et al., 2006), as well as in reward-related behaviors and feeding (Gallagher et al., 1990; Kentridge et al., 1991; Kim et al., 2017; Robinson et al., 2014; Seo et al., 2016). For instance, previous studies showed that the somatostatin-expressing (SOM^+^) subpopulation of neurons in the CeL is essential for the acquisition and expression of conditioned freezing and action suppression (Fadok et al., 2017; Li et al., 2013; Yu et al., 2016). The protein kinase C-δ- expressing (PKC-δ^+^) CeL neurons, on the other hand, participate in conveying aversive US information, drive avoidance response, and are sufficient to instruct aversive learning (Yu et al., 2017). Furthermore, it has been shown that CeA neurons, including PKC-δ^+^ CeL neurons, are critical for suppression of feeding behavior (Cai et al., 2014; Petrovich et al., 2009). As the CeL is a major direct downstream target of the IC, these findings point to the possibility that the IC controls learning or expression of taste-motivated behavioral inhibition through the IC-CeL circuit.

To test this hypothesis, we used anatomical, electrophysiological, and circuit based manipulation approaches. We found that an excitatory monosynaptic connection exists between the posterior division of the IC and the CeL in mice, and that activation of the IC-CeL pathway excites specific subtypes of neurons within the CeL. Notably, selective inhibition of CeL-projecting IC neurons specifically impairs the conditioned inhibitory response to a cue predicting an aversive tastant. Furthermore, activation of the IC-CeL circuit with optogenetics produces a powerful suppression of ongoing licking behavior in thirsty mice, induces avoidance behavior, and is sufficient to instruct conditioned lick suppression. These results reveal an important role of the IC-CeL circuit in the establishment of anticipatory behavioral inhibition, in particular the inhibition of consummatory behavior in response to cues predicting an unpleasant taste.

## Materials and Methods

### Animals

Before surgery, mice were group-housed under a 12-h light-dark cycle (7 a.m. to 7 p.m. light) with food and water freely available. The *Som-cre* (Taniguchi et al., 2011), *Ai14* (Madisen et al., 2010), *Prkcd-cre* (Haubensak et al., 2010), and *Rosa26-stop^flox^-tTA* (Li et al., 2010) mice were described previously and were purchased from the Jackson Laboratory. All mice were bred onto C57BL/6J genetic background. The *Som-cre;Ai14* mice, which were heterozygous for both the Cre allele and the Lox-Stop-Lox-tdTomato allele, were bred by crossing homozygous *Som-cre* mice with homozygous *Ai14* reporter mice. Male mice of 40–80 d of age were used for behavioral and anatomical experiments. Male and female mice of 35-45 d of age were used for *in vitro* slice physiology experiments. Behavioral experiments were performed during the light cycle. All procedures involving animals were approved by the Institute Animal Care and Use Committees of Cold Spring Harbor Laboratory and carried out in accordance with US National Institutes of Health standards.

### Viral vectors

Most of the adeno-associated viruses (AAV) were produced by the University of North Carolina vector core facility or the University of Pennsylvania vector core and have previously been described (Ahrens et al., 2015; Penzo et al., 2015; Stephenson-Jones et al., 2016; Yu et al., 2016): AAV9-Ef1a-DIO-eYFP, AAV9-Ef1a-DIO-hChR2(H134R)-eYFP, AAV9-CAG-ChR2-GFP, AAV9.CAG.Flex.TeLC-eGFP.WPRE.bGH, and AAV-TRE-hGFP-TVA-G. The AAV8.2-hEF1α-DIO-synaptophysin-mCherry was produced by the MIT Viral Gene Transfer Core. The EnvA-pseudotyped, protein-G-deleted rabies-EnvA-SAD-ΔG-mCherry virus was produced by the Viral Vector Core Facility at Salk Institute (Penzo et al., 2015). CAV2-Cre was purchased from Montpellier vector platform (Plateforme de Vectorologie de Montpellier (PVM), Biocampus Montpellier, Montpellier, France) (Penzo et al., 2015; Stephenson-Jones et al., 2016). All viral vectors were stored in aliquots at −80 °C until use.

### Histology

Animals were deeply anesthetized and transcardially perfused with PBS, followed by perfusion with 4% paraformaldehyde (PFA) in PBS. Brains were dissected out and postfixed in 4% PFA at 4°C for three hours followed by cryoprotection in a PBS-buffered sucrose (30%) solution until brains were saturated (~36 h). 50 μm coronal brain sections were cut on a freezing microtome (SM 2010R, Leica). Brain sections were first washed in PBS (3 x 5 min) at room temperature (RT) and then were blocked in 3% normal goat serum (NGS) in PBST (0.3% Triton X-100) for 30 min at RT, followed by incubation with primary antibodies overnight at 4 °C. Sections were then washed with PBS (4 x 15 min) and incubated with fluorescent secondary antibodies at RT for 2 hours. After washing with PBS (4 x 15 min), sections were mounted onto glass slides with Fluoromount-G (Beckman Coulter). Images were taken using a LSM 780 laser-scanning confocal microscope (Carl Zeiss).

### Electrophysiology

For electrophysiological experiments, mice were anesthetized with isoflurane, decapitated and their brains quickly removed and chilled in ice-cold dissection buffer (110.0 mM choline chloride, 25.0 mM NaHCO_3_, 1.25 mM NaH_2_PO_4_, 2.5 mM KCl, 0.5 mM CaCl_2_, 7.0 mM MgCl_2_, 25.0 mM glucose, 11.6 mM ascorbic acid and 3.1mM pyruvic acid, gassed with 95% O_2_ and 5% CO_2_). Coronal slices (300 μm) containing the amygdala complex were cut in dissection buffer using a HM650 Vibrating-blade Microtome (Thermo Fisher Scientific). Slices were immediately transferred to a storage chamber containing artificial cerebrospinal fluid (ACSF) (118 mM NaCl, 2.5 mM KCl, 26.2 mM NaHCO_3_, 1 mM NaH_2_PO_4_, 20 mM glucose, 2 mM MgCl_2_ and 2 mM CaCl_2_, at 34 °C, pH 7.4, gassed with 95% O_2_ and 5% CO_2_). After 40 min recovery time, slices were transferred to RT (20–24°C) and perfused with ACSF constantly.

Simultaneous whole-cell patch-clamp recordings from pairs of SOM^+^ and SOM^−^ CeL neurons were obtained with Multiclamp 700B amplifiers (Molecular Devices). Recordings were under visual guidance using an Olympus BX51 microscope equipped with both transmitted light illumination and epifluorescence illumination, and SOM^+^ cells were identified based on their fluorescence (tdTomato). To evoke IC-driven synaptic transmission onto CeL neurons, the AAV-ChR2-YFP was injected into the IC of *Som-Cre;Ai14* mice and allowed to express for 3 weeks. Acute brain slices were prepared, and a blue light was used to stimulate ChR2-expressing axons in the CeL. The light source was a single-wavelength LED system (λ = 470 nm; CoolLED.com) connected to the epifluorescence port of the Olympus BX51 microscope. 1 ms light pulses were triggered by a TTL signal from the Clampex software to drive synaptic responses. Light pulses were delivered every 10 seconds and synaptic responses were low-pass filtered at 1 KHz and recorded at holding potentials of −70 mV (for AMPA-receptor-mediated responses) and +40 mV (for NMDA-receptor-mediated responses). NMDA-receptor-mediated responses were quantified as the mean current amplitude from 50-60 ms after stimulation. Evoked EPSCs were recorded in ACSF with 100 μM picrotoxin to block inhibitory synaptic transmission. The internal solution for voltage-clamp experiments contained 115 mM cesium methanesulphonate, 20 mM CsCl, 10 mM HEPES, 2.5 mM MgCl_2_, 4 mM Na_2_-ATP, 0.4 mM Na_3_GTP, 10 mM Na-phosphocreatine and 0.6 mM EGTA (pH 7.2). To assess presynaptic function, a paired-pulse stimulation protocol (50 ms inter-stimulus interval) was used to evoke double-EPSCs, and paired-pulse ratio (PPR) was quantified as the ratio of the peak amplitude of the second EPSC to that of the first EPSC.

### Monosynaptic tracing with pseudotyped rabies virus

Retrograde tracing of monosynaptic inputs onto genetically-defined cell populations of CeL was performed and described in our previous study (Penzo et al., 2015), and the data presented here were generated from the same study but were not published previously. Briefly, the *Som-cre;Rosa26-stop^flox^-tTA* mice and the *Prkcd-cre;Rosa26-stop^flox^-tTA* mice, which express tTA in SOM^+^ cells and PKC-δ^+^ cells, respectively, were injected into the CeL with the AAV-TRE-hGFP-TVA-G (0.2–0.3 μl) that expresses the following components in a tTA-dependent manner: a fluorescent reporter histone GFP (hGFP); TVA (which is a receptor for the avian virus envelope protein EnvA); and the rabies envelope glycoprotein (G). Two weeks later the mice were injected in the same location with the rabies-EnvA-SAD-ΔG-mCherry (1.2 μl), a rabies virus that is pseudotyped with EnvA, lacks the envelope glycoprotein, and expresses mCherry. This method ensures that the rabies virus exclusively infects cells expressing TVA. Furthermore, complementation of the modified rabies virus with envelope glycoprotein in the TVA-expressing cells allows the generation of infectious particles, which then can trans-synaptically infect presynaptic neurons.

### Stereotaxic surgery

Standard surgical procedures were followed for stereotaxic injection (Li et al., 2013; Penzo et al., 2015). Briefly, mice were anesthetized with ketamine (100 mg per kg of body weight) supplemented with dexmedetomidine hydrochloride (0.4 mg per kg) and positioned in a stereotaxic injection frame (myNeuroLab.com). A digital mouse brain atlas was linked to the injection frame to guide the identification and targeting (Angle Two Stereotaxic System, myNeuroLab.com).

Viruses (~0.4 μl) were delivered with a glass micropipette (tip diameter, ~5 μm) through a skull window (1–2 mm^2^) by pressure applications (5–20 psi, 5–20 ms at 0.5 Hz) controlled by a Picrospritzer III (General Valve) and a pulse generator (Agilent). The injection was performed at the following stereotaxic coordinates for CeL: 1.18 mm posterior, 2.9 mm lateral, and 4.6 mm ventral from bregma; and for IC: 0.10 mm posterior, 3.90 mm lateral, and 4.20 mm ventral from bregma. For optogenetic experiments, immediately after viral injection, an optical fiber (core diameter, 105 μm; Thorlabs, Catalog number FG105UCA) was implanted 300 μm above the center of viral injection. The optical fiber together with the ferrule (Thorlabs) was secured to the skull with C&B-Metabond Quick adhesive luting cement (Parkell Prod), followed by dental cement (Lang Dental Manufacturing).

Following the above procedures, a small piece of metal bar was mounted on the skull, which was used to hold the mouse in the head fixation frame during behavior experiments.

### Behavioral tasks

#### Licking Behavior

Water deprivation started 23 hours before training. Mice were trained in the head fixation frame for 10 minutes daily. A metal spout was placed in front of the animal’s mouth for water delivery. The spout also served as part of a custom “lickometer” circuit, which registered a lick event each time a mouse completed the circuit by licking the spout while standing on a metal floor. The lick events were recorded by a computer through custom software written in LabView (National Instruments). Each lick triggered a single opening of a water valve calibrated to deliver 0.3 μl water.

It took mice 4–7 days to achieve stable licking, the criterion for which was 10-minute continuous licking with no interval between licking longer than 10 seconds. We used a lick suppression index to quantify animals’ degree of photostimulation-evoked suppression of licking behavior: Lick suppression index = (L_PRE_ – L_CS_) / (L_PRE_ + L_CS_), where L_PRE_ is the number of licks in the 5 s period before CS onset, and L_CS_ is the number of licks in the 5 s CS period (Yu et al., 2016).

#### Go/no-go task

Water deprivation started 23 hours before training, and mice were habituated to the head fixation frame for 20 minutes on the first day of training with access to water through the metal spout. On following days, animals underwent 2 training sessions each day, one in the morning and the other in the afternoon. The 2 sessions were at least 4 hours apart, with each consisting of 100 trials. For the subsequent 3-6 sessions, mice were exposed only to the “go” cue (a 1-s, 5-kHz pure tone) followed by the delivery of 4.5 μl of water. After mice successfully retrieved water on at least 80% of the trials, they moved to the next training phase, in which they were required to lick the spout at least 1 time during the go cue in order for the water to be released. This phase took an additional 3-6 sessions until the animals reached the criteria of 80% correct responses. Following this phase, animals received 1 training session consisting of the go cue paired with the delivery of sucrose solution (100 mM) instead of water.

The next phase consisted of 10 sessions of go/no-go training. During this phase, 50 presentations of the go cue were delivered randomly intermixed with 50 presentations of the “no-go” cue (a 1-s white noise), with the constraint that either cue could not appear more than 5 times in a row, and that the first trial was always a go cue. Licking the spout during the no-go cue resulted in the delivery of quinine solution (4.5 μl, 5 mM). The mice were required to lick at least once the spout during the 1 s window of cue presentation in order to receive the US. During all phases of the experiment, brief suction (500 ms in duration) near the spout was applied 3.5 s after tone onset to remove any residual solution from the previous trial.

For analysis, trials were sorted into go trials and no-go trials. A correct response during a go trial (“hit”) occurred when the mouse successfully licked the spout during the go cue and subsequently received sucrose. A correct response during a no-go trial (“correct reject”) occurred when the mouse successfully omitted the lick response during the no-go cue and thus avoided quinine. The overall performance over the entire session was calculated as the total correct responses divided by the total trials: overall performance = (hits + correct rejects) / (total trials).

For the optogenetics experiments we used a modified version of this go/no-go task, in which licking during the go cue led to water delivery (4.5 μl), whereas licking during the no-go cue resulted in water delivery (4.5 μl) accompanied by laser stimulation. The laser was delivered coincidentally with water delivery (50 ms after CS offset) at 20- or 30-Hz for 2.5 seconds (the duration that the water would be available if the animal licked during the no-go cue presentation). Suction was applied to remove any unconsumed water. Animals received 8 training sessions in the final phase of this task.

#### Real time place aversion (RTPA)

As previously described (Stephenson-Jones et al., 2016), one side of a custom chamber (23 × 33 × 25 cm; made from plexiglass) was assigned as the stimulation zone, counterbalanced among mice. Mice were placed individually in the middle of the chamber at the onset of the experiment, the duration of which was 30 min. Laser stimulation (5-ms pulses delivered at 20 or 30 Hz) was triggered when mice entered the stimulation zone, and lasted until mice exited the stimulation zone. Mice were videotaped with a CCD camera interfaced with the Ethovision software (Noldus Information Technologies), which was used to control the laser stimulation and extract the behavioral parameters (position, time, distance, and velocity).

### In vivo optogenetics

For bilateral optogenetic stimulation in the CeL, a branched patch-cord (Doric Lenses, Catalog number BFP(2)_105/125/900-0.22_1m_FC-2xZF1.25) for light delivery was connected at one end to a laser source (λ = 473 nm, OEM Laser Systems) and at the other end, which was composed of two terminals, to two CeL-implanted optical fibers through sleeves (Thorlabs). For photostimulation-induced lick suppression, the stimuli were 5-ms 30-Hz light pulses (or across a range of frequencies) delivered for 5 s. For photostimulation during RTPA, 5-ms 20- or 30-Hz light pulses were delivered. Laser intensity was 10 mW measured at the end of optical fiber.

### Statistics and data presentation

All data are presented as mean ± s.e.m. All statistics are indicated where used. Data were analyzed with GraphPad Prism. Behavioral tests were performed by an investigator with knowledge of the identity of the experimental groups. All behavior experiments were controlled by computer systems, and data were collected and analyzed in an automated and unbiased way. Virus-injected animals in which the injection or optical fiber implantation was misplaced were excluded.

## Results

To investigate the function of the IC-CeL circuit, we began by characterizing how IC neurons innervate the major CeL cell types. We first used a modified rabies virus system to trace the monosynaptic inputs onto CeL neurons (Callaway and Luo, 2015; Penzo et al., 2015). This approach revealed a monosynaptic projection from the IC to the CeL (Fig. 1A, B). In particular, IC neurons innervate both of the two major populations of the CeL, the SOM^+^ neurons and PKC-δ^+^ neurons (Fig. 1A, B). Notably, the IC was the only cortical region identified by this approach to send monosynaptic projections to the CeL. The CeL-projecting IC neurons were preferentially localized in the posterior part of the IC, which overlaps at least partially with the GC (Fig. 1B).

**Figure 1.**
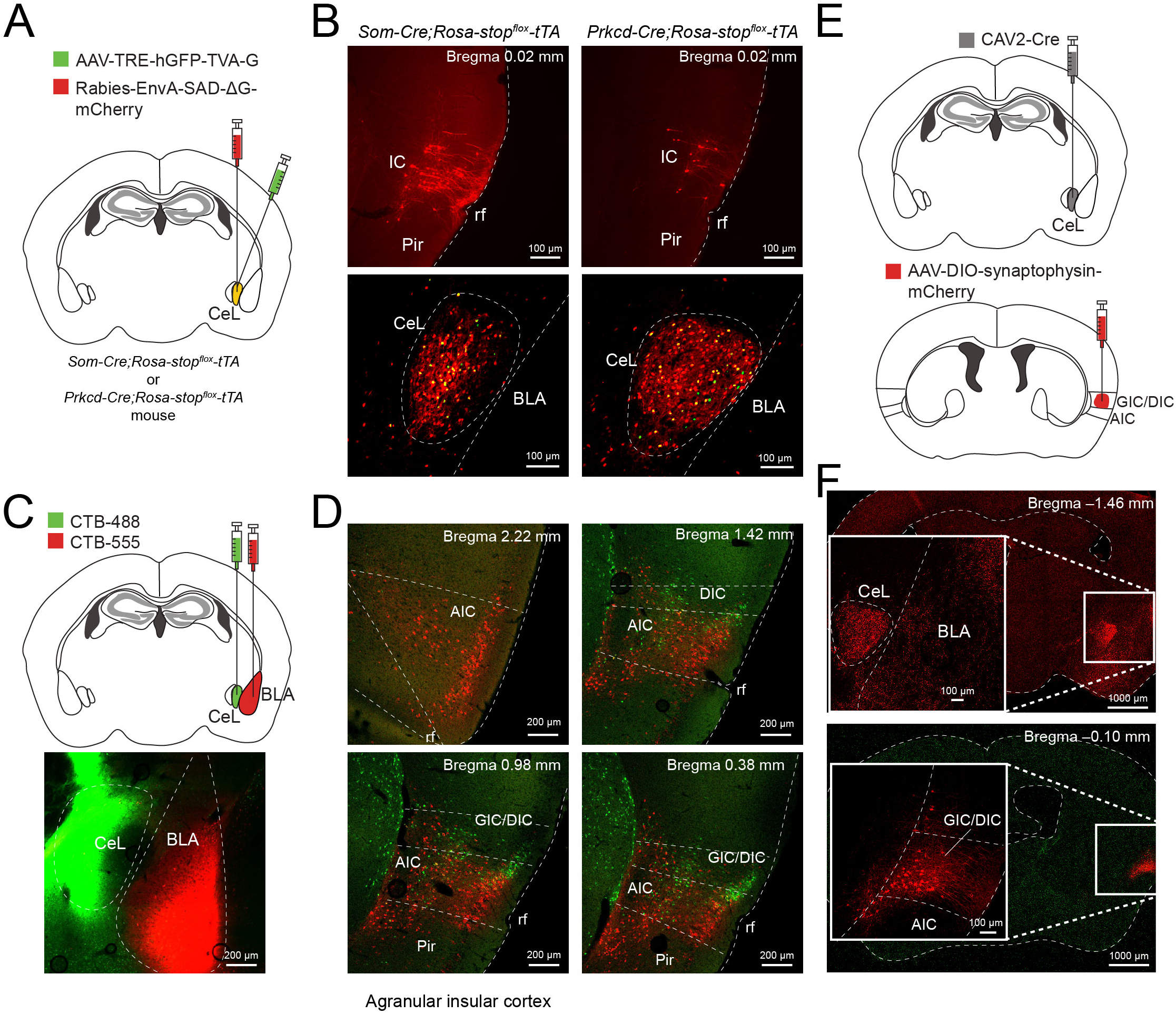
The IC sends monosynaptic projections to the amygdala. (**A**) A schematic of the experimental approach (see Methods). (**B**) Representative images of the tracing result for SOM^+^ (left) and PKC-δ^+^ (right) CeL neurons. Top, retrogradely labeled neurons in the IC. Bottom, starter neurons in the CeL are identified by their co-expression of mCherry and histone GFP (cells in yellow). Data in A & B were replicated in 3 mice for each group, and were from the same injections reported in (Penzo et al., 2015). (**C**) Top: a schematic of the experimental approach. Bottom: a representative image of the injection sites. (**D**) The distribution of the BLA-projecting (red) and the CeL-projecting (green) neurons in the IC. AIC, agranular insular cortex; GIC, granular insular cortex; DIC, disgranular insular cortex. Data in C & D were replicated in two mice. (**E**) A schematic of the experimental approach. (**F**) Representative images showing the mCherry^+^ axons (top), which originated from the CeL-projecting IC neurons (bottom). In the insets are enlarged images of the boxed areas in the right, which are located in the amygdala (top) and the IC (bottom).

As the IC also sends projections to the BLA (Allen et al., 1991), we next determined whether the same IC neurons project to both the CeL and the BLA. To this end we injected the CeL and BLA with the retrograde tracer AlexaFluor-488- or 555-conjugated cholera toxin (CTB-488 or CTB-555), respectively (Fig. 1C). We found that CTB reliably labeled CeL-projecting IC neurons (Fig. 1D), with a distribution pattern similar to that of CeL-projecting IC neurons labeled by the rabies virus (Fig. 1B). CTB also labeled a prominent population of BLA-projecting IC neurons, the vast majority of which has a distribution pattern distinct from that of CeL-projecting IC neurons (Fig. 1D). This result demonstrates that CeL-projecting neurons and BLA-projecting neurons in the IC are largely non-overlapping populations.

To selectively target the CeL-projecting IC neurons and visualize their axonal projections, we injected the CeL with a retrograde canine adenovirus expressing Cre recombinase (CAV2-Cre) (Bru et al., 2010), followed by injecting the IC with an adeno-associated virus harboring a double-floxed inverted open reading frame (AAV-DIO) that expresses, in a Cre-dependent manner, a presynaptic protein synaptophysin tagged with a fluorescent protein mCherry (AAV-DIO-synaptophysin-mCherry) (Fig. 1E). This strategy led to the labeling of IC neurons that sent dense axonal fibers to the CeL (Fig. 1F). Sparse labeling of axon fibers in the BLA can also be detected (Fig. 1F), which likely was caused by spillover of the CAV2-Cre to the BLA and thus the labeling of BLA-projecting IC neurons. Though the ipsilateral CeL had the densest projections, we also observed mCherry^+^ axon terminals in the contralateral CeL, consistent with results from retrograde tracing with CTB (data not shown). Altogether, the anatomical tracing results demonstrate that the IC sends robust projections to the CeL, and that the IC-CeL and IC-BLA are distinct circuits.

To assess the synaptic connectivity between the IC and CeL, we used the *Som-cre;Ai14* mice, in which SOM^+^ cells can be identified by their red fluorescence, and injected the IC of these mice with an AAV expressing the light-gated cation channel channelrodhopsin-2 (AAV-ChR2-YFP) that allows photostimulation of axonal projections (Zhang et al., 2006) (Fig. 2A). Approximately 3 weeks after the AAV injection, we prepared from these mice acute brain slices containing the CeL, in which we recorded synaptic transmission onto simultaneously patched pairs of adjacent SOM^+^ (red-fluorescent) and SOM^−^ (non-fluorescent) CeL neurons in response to optogenetic stimulation of the IC inputs. Brief light pulses evoked fast excitatory synaptic transmission in nearly all the recorded CeL neurons (Fig. 2B). Notably, the AMPA receptor-mediated component of synaptic transmission onto SOM^+^ neurons was significantly greater than that onto SOM^−^ neurons (Fig. 2B-C). The NMDA receptor-mediated synaptic transmission onto these two neuronal populations was not different. In a subset of these pairs, we also examined paired-pulse ratio (PPR; see Methods) and found no significant difference between the two cell types (Fig. 2C). These results indicate that IC inputs can activate both SOM^+^ neurons and SOM^−^ neurons in the CeL, with the latter being mainly PKC-δ^+^ neurons (Li et al., 2013).

**Figure 2:**
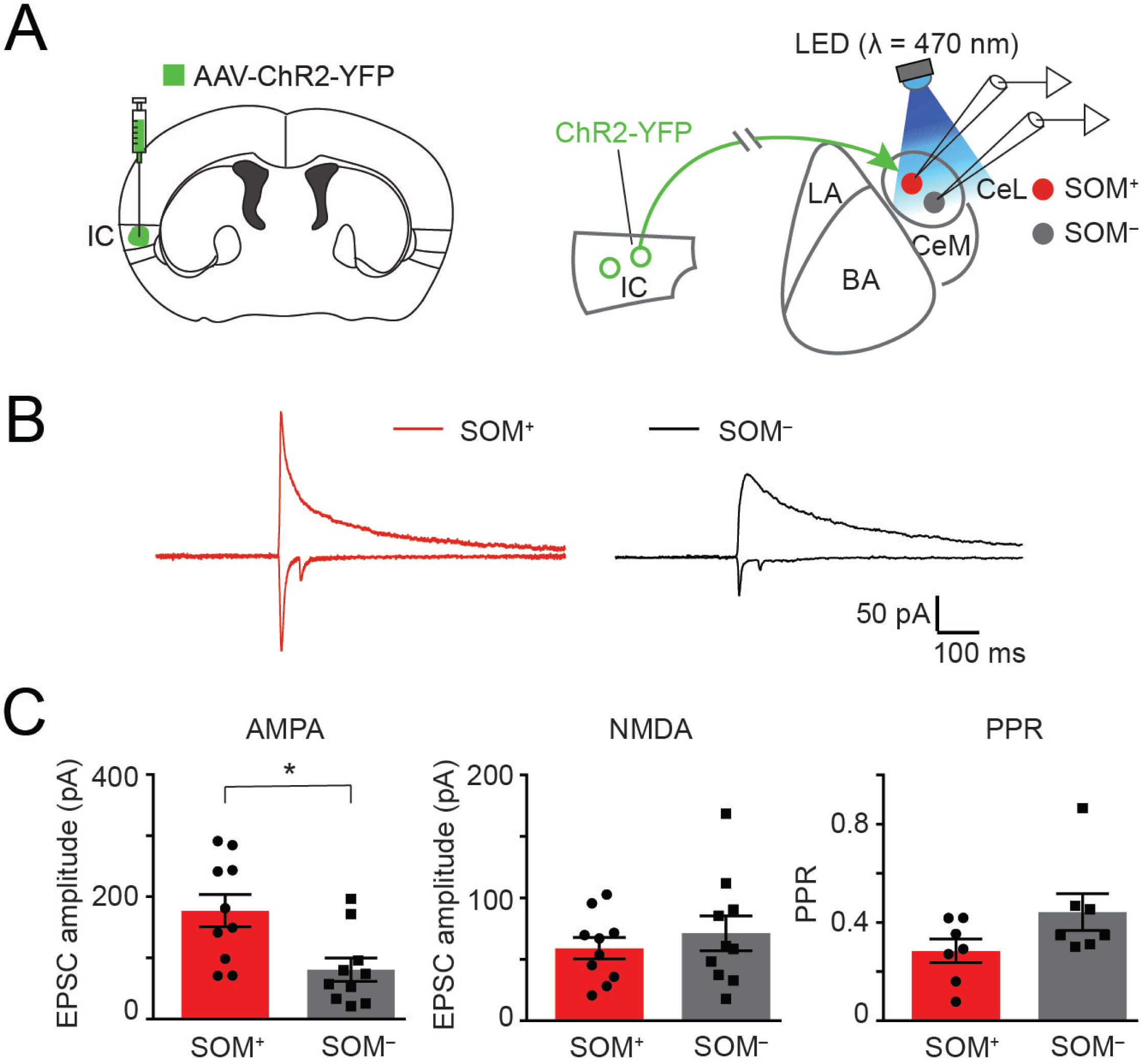
Functional connectivity between IC neurons and CeL neurons. (**A**) A schematic of the experimental design. AAV-ChR2-YFP was injected into the IC of a *SOM;Ai14 mouse* (left), and patch clamp recording was performed in acute slices containing the amygdala (right). (**B**) Sample traces of EPSCs recorded from a SOM^+^ and a SOM^−^ CeL neuron, which were recorded simultaneously. The EPSCs were evoked by optogenetic stimulation of the IC axons terminating in the CeL, with the AMPA receptor-mediated EPSCs being stimulated with two pulses (50 ms inter-pulse interval) protocol. (**C**) Quantification of the EPSC amplitude evoked by the first pulse mediated by AMPA receptors is shown in the left panel, NMDA receptors is shown in the middle, and the paired-pulse ratio (PPR) of AMPA receptor-mediated EPSCs is shown on the right. T-tests revealed that the AMPA receptor-mediated EPSC was larger in SOM^+^ than SOM-CeL neurons (AMPA, t(18) = 3, \**P* = 0.0085, NMDA, t(18) = 0.7, *P* = 0.47, n = 10 pairs, *t* test; PPR, t(12) = 1.8, *P* = 0.1, n = 7 pairs, *t* test). Data in C are presented as mean ± s.e.m.

We reasoned that the IC-CeL circuit could have a role in behavioral inhibition, because SOM^+^ CeL neurons are essential for the generation of passive defensive responses, including freezing behavior and action suppression (Fadok et al., 2017; Li et al., 2013; Penzo et al., 2015; Yu et al., 2016), while PKC-δ^+^ CeL neurons convey aversive US information and are sufficient to instruct aversive learning (Yu et al., 2017). To test this hypothesis, we set out to inhibit the CeL-projecting IC neurons in mice and subsequently trained these mice in a go/no-go task that engages the IC. We first injected the CeL with the CAV2-Cre (Fig. 3A), and then injected the IC in the same mice with an AAV expressing the tetanus toxin light chain (TeLC), which blocks neurotransmitter release (Murray et al., 2011), or GFP (as a control) in a Cre-dependent manner (AAV-DIO-TeLC-GFP or AAV-DIO-GFP, respectively) (Fig. 3A). As described above (Fig. 1E, F), this strategy led to selective targeting of the IC-CeL circuit (Fig. 3B, C).

**Figure 3.**
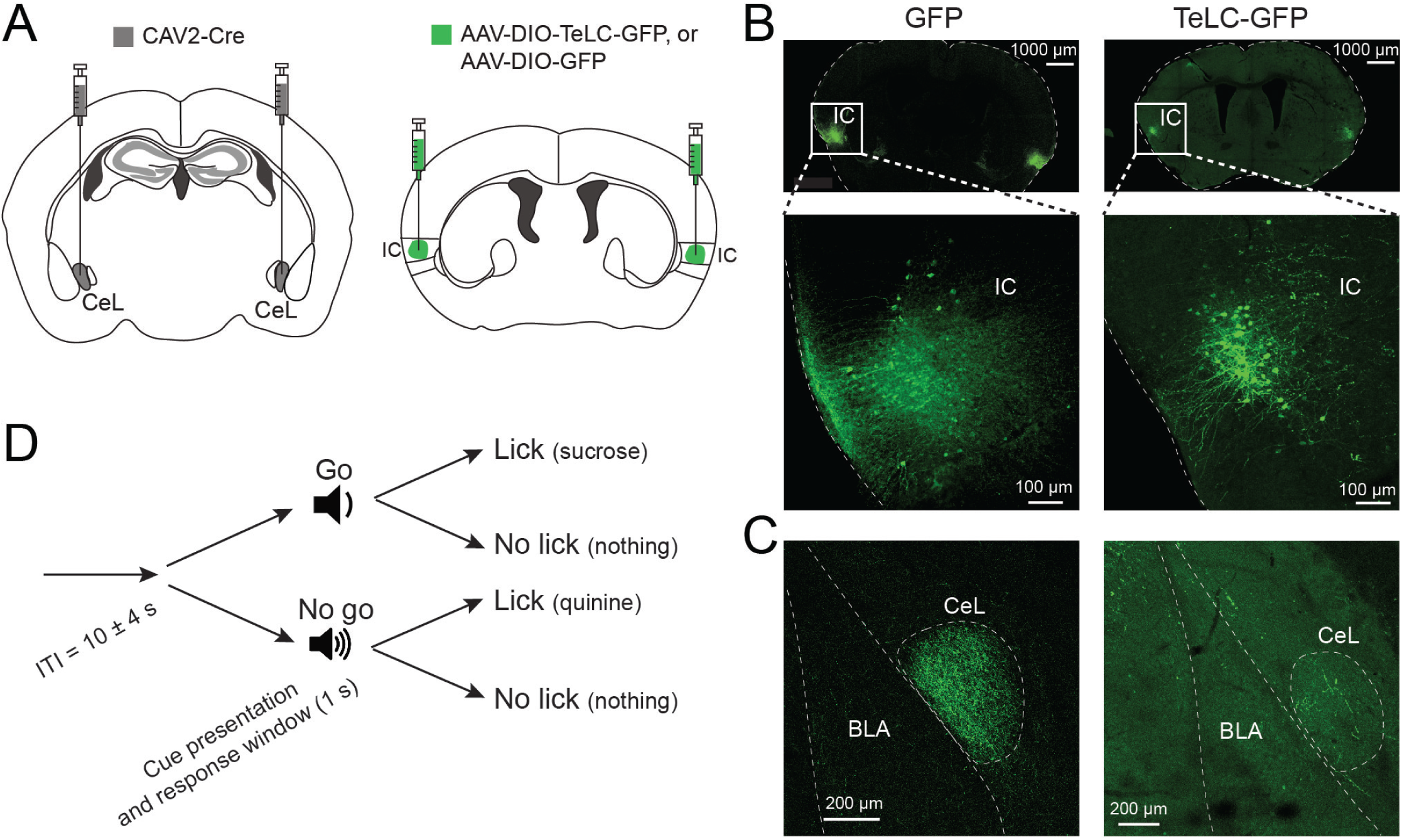
Experimental design to test the role of the IC-CeL circuit in a go/no-go task. (**A**) A schematic of the experimental design to selectively inhibit the IC-CeL circuit. (**B**) Representative images showing the cells in the IC infected with GFP (left panels) and TeLC-GFP (right panels) virus. In the lower panel are enlarged images of the boxed areas in the images in the upper panel. (**C**) Axon terminals expressing GFP (left) or TeLC-GFP (right), which originated from CeL-projecting IC neurons. (**D**) A schematic of the go/no-go task.

Four to five weeks following viral injections, we began training these mice in the go/no-go task (see Methods), in which an auditory stimulus (go cue) predicts the delivery of a palatable liquid (sucrose), while a different auditory stimulus (no-go cue) predicts the delivery of an unpleasant liquid (quinine) (Fig. 3D). Mice need to learn to produce an instrumental response (lick) during the go cue to receive sucrose, and inhibit that response during the no-go cue to avoid quinine (Fig. 4A). Learning in this task has previously been shown to be paralleled by the development of cue-specific responses in IC neurons (Gardner and Fontanini, 2014).

**Figure 4.**
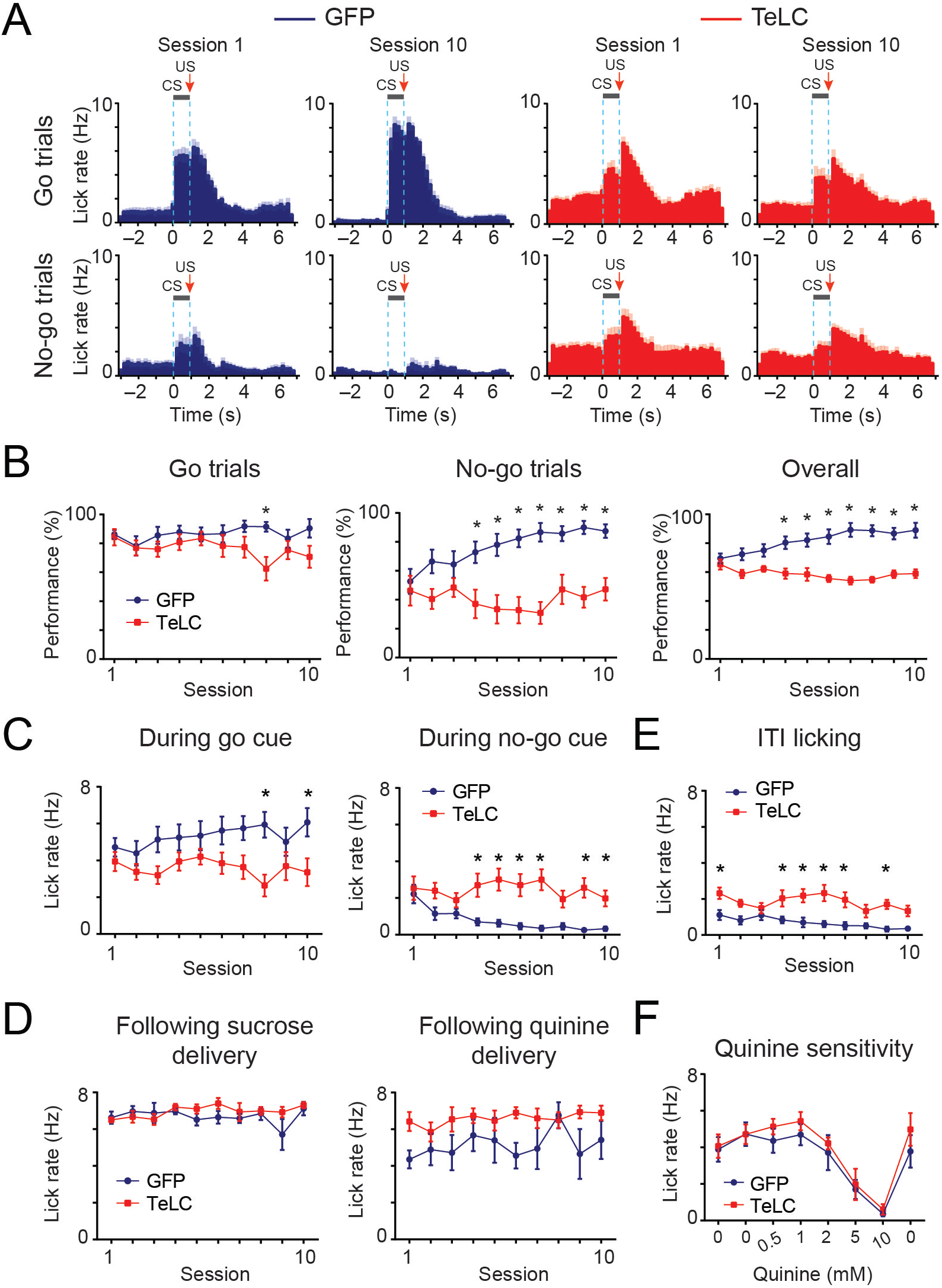
The IC-CeL circuit is required for suppression of behavioral responding. (**A**) Population histogram of licking behavior during the first and final sessions of go/no-go training. Licking behavior is represented separately according to go (top panels) or no-go (bottom panels) trials for both GFP (blue; n = 9) and TeLC (red; n = 7) mice. Dashed line denotes the time window of CS delivery. US was delivered 50 ms after CS offset. Shaded regions represent s.e.m. (**B**) Quantification of performance (the percentage of correct responding trials) on the go/no-go task. For the go trials, the TeLC animals showed a mild reduction in performance compared to GFP animals (two-way repeated measures (RM) ANOVA revealed a significant interaction, but no main effects: main effect of session, F(9,126) = 1.4; *P* = 0.2; main effect of virus treatment, F(1,14) = 2.4; *P* = 0.1; interaction, F(9,126) = 2.8; *P* < 0.01; \**P* < 0.01, Post hoc Sidak’s multiple comparisons test). For the no-go trials, the TeLC animals showed a marked reduction in performance compared to GFP animals (two-way RM ANOVA revealed a main effect of session (F(9,126) = 2.84; *P* = 0.005), a main effect of virus treatment (F(1,14) = 17.56; *P* < 0.001), and a significant interaction (F(9,126) = 4.8; *P* < 0.001); \**P* < 0.05, Post hoc Sidak’s multiple comparisons test). For the overall performance, the TeLC animals showed a reduction in performance compared to GFP animals (two-way RM ANOVA revealed a main effect of session (F(9,126) = 2.41; *P* = 0.01), a main effect of virus treatment (F(1,14) = 24.28; *P* < 0.01), and a significant interaction (F(9,126) = 9.8; *P* < 0.01). \**P* < 0.01, Post hoc Sidak’s multiple comparisons test). (**C**) TeLC animals showed a mild reduction in licking in response to the go cue (two-way RM ANOVA revealed a significant interaction, but no main effects (main effect of session, F(9,126) = 1.9; *P* = 0.06; main effect of virus treatment, F(1,14) = 4.1; *P* = 0.06; interaction, F(9,126) = 3.5; *P* < 0.001; \**P* < 0.05, Post hoc Sidak’s multiple comparisons test). TeLC animals showed an increase in licking in response to the no-go cue (two-way RM ANOVA revealed a main effect of session (F(9,126) = 3.76; *P* < 0.001), a main effect of virus treatment (F(1,14) = 15.8; *P* = 0.001), and a significant interaction (F(9,126) = 4.45; *P* < 0.001; \**P* < 0.05, Post hoc Sidak’s multiple comparisons test). (**D**) TeLC inhibition of the IC-CeL circuit did not affect US-evoked licking rate (*P* > 0.05, two-way RM ANOVA). (**E**) TeLC inhibition of the IC-CeL circuit increased the inter-trial licking rate (two-way RM ANOVA revealed a main effect of session (F(9,126) = 4.13; *P* < 0.001), a main effect of virus treatment (F(1,14) = 15.73; *P* = 0.001), and a significant interaction (F(9,126) = 2.08; *P* = 0.037); \**P* < 0.05, Post hoc Sidak’s multiple comparisons test). (**F**) Quinine sensitivity test showing lick rate during a 10-minute free-licking session for various concentrations of quinine followed by a final session of water. TeLC inhibition of the IC-CeL circuit did not affect quinine sensitivity (two-way RM ANOVA main effect of quinine concentration, F(7,84) = 16.3, *P* < 0.0001; main effect of virus, F(1,12) = 0.78, *P* = 0.4; interaction, F(7,84) = 0.24, *P* = 0.97). All data are presented as mean ± s.e.m.

We found that bilateral inhibition of synaptic transmission from CeL-projecting IC neurons with TeLC markedly affected animals’ behavior in the no-go trials, but left that in the go trials of this task largely unaffected (Fig. 4A, B). Specifically, in the go trials, mice in both the GFP group and the TeLC group showed stimulus-evoked licking (Fig. 4A), leading to similar performance (Fig. 4B); although closer inspection of these animals’ behavioral patterns revealed that the TeLC mice did not allocate their licking to the CS and US period as much as the GFP control mice, especially on the final training session (Fig. 4A,B). By contrast, in the no-go trials, while the GFP mice gradually learned to withhold licking in response to the no-go cue, and thus successfully avoid quinine in most of the trials towards the end of the training sessions (Fig. 4A, B), the TeLC mice showed no sign of learning and thus were markedly impaired in performance even at the end of the training sessions (Fig. 4A, B). The overall performance of the GFP mice also showed a learning effect, evidenced by a gradual increase in performance over the first 8 training sessions followed by asymptotic performance (Fig. 4B). On the other hand, the animals expressing TeLC showed no such improvement with continued training (Fig. 4B).

The impairment in behavioral inhibition observed during the no-go trials (or decrease in the “correct reject”) in the TeLC mice could be caused by a general increase in responding. If so, then the performance of these mice in the go trials (measured as the percentage of trials in which the mice made a response; or the “hit” rate) would also increase. However we observed no such increase; in fact, the TeLC group showed a mild decrease in hit rate, in particular in late sessions (Fig. 4B). The TeLC mice also showed a reduced lick rate in response to the go cue, and had lick rate similar to that of the GFP mice following the delivery of sucrose (Fig. 4A, C, D), further arguing against a general increase in responding in these mice. In contrast, these mice showed an increased lick rate specifically to the no-go cue and during the inter-trial interval (ITI) (Fig. 4A, C, E), consistent with the notion that they were impaired in action suppression. We noticed that the TeLC mice had a tendency to show increased lick rate following quinine delivery compared with the GFP mice (Fig. 4D), suggesting that inhibiting the IC-CeL pathway may partially impair the processing of aversive taste information during this task. Alternatively, or additionally, the observations that the TeLC mice showed impaired performance (although mild) and reduced lick rate during the go cue (Fig. 4B,C), as well as impaired performance and increased lick rate during the no-go cue and quinine delivery (Fig. 4B-D) could be explained by an impairment in these mice in anticipation of salient outcomes, a function that has been attributed to the CeA (Balleine and Killcross, 2006; Haney et al., 2010; Roesch et al., 2012).

We also tested a subset of these animals on their sensitivity to increasing concentrations of quinine during a free-licking session (10 minute). Similar to the GFP control group, the TeLC group showed decreased average licking rate during this period (reflecting a reduction in the total volume consumed) with increasing concentrations of quinine (Fig. 4F). This result is consistent with previous findings that lesions of the GC in rats do not affect the amount of either quinine or sucrose solutions consumed at varying concentrations (Hashimoto and Spector, 2014), and indicates that inhibition of CeL-projecting IC neurons does not abolish animals’ basic ability to process quinine’s sensory and aversive properties, at least when there is little cognitive demand. Together, these results indicate that the IC-CeL pathway is required for establishing the learned, anticipatory behavioral inhibition to avoid an aversive tastant.

Our results suggest that the CeL-projecting IC neurons preferentially regulate the no-go response, consistent with the critical role of the CeL in processing aversive information and in the learning and expression of avoidance behaviors. We therefore tested whether activation of the IC-CeL pathway is sufficient to drive aversive responses as well as instruct learning of an avoidance behavior. For this purpose we delivered the ChR2 or GFP (as a control) specifically into CeL-projecting IC neurons bilaterally, using the retrograde and intersectional strategy based on the CAV2-Cre as described above (Fig. 5A), and subsequently bilaterally implanted optical fibers over the CeL (Fig. 5B, C).

**Figure 5.**
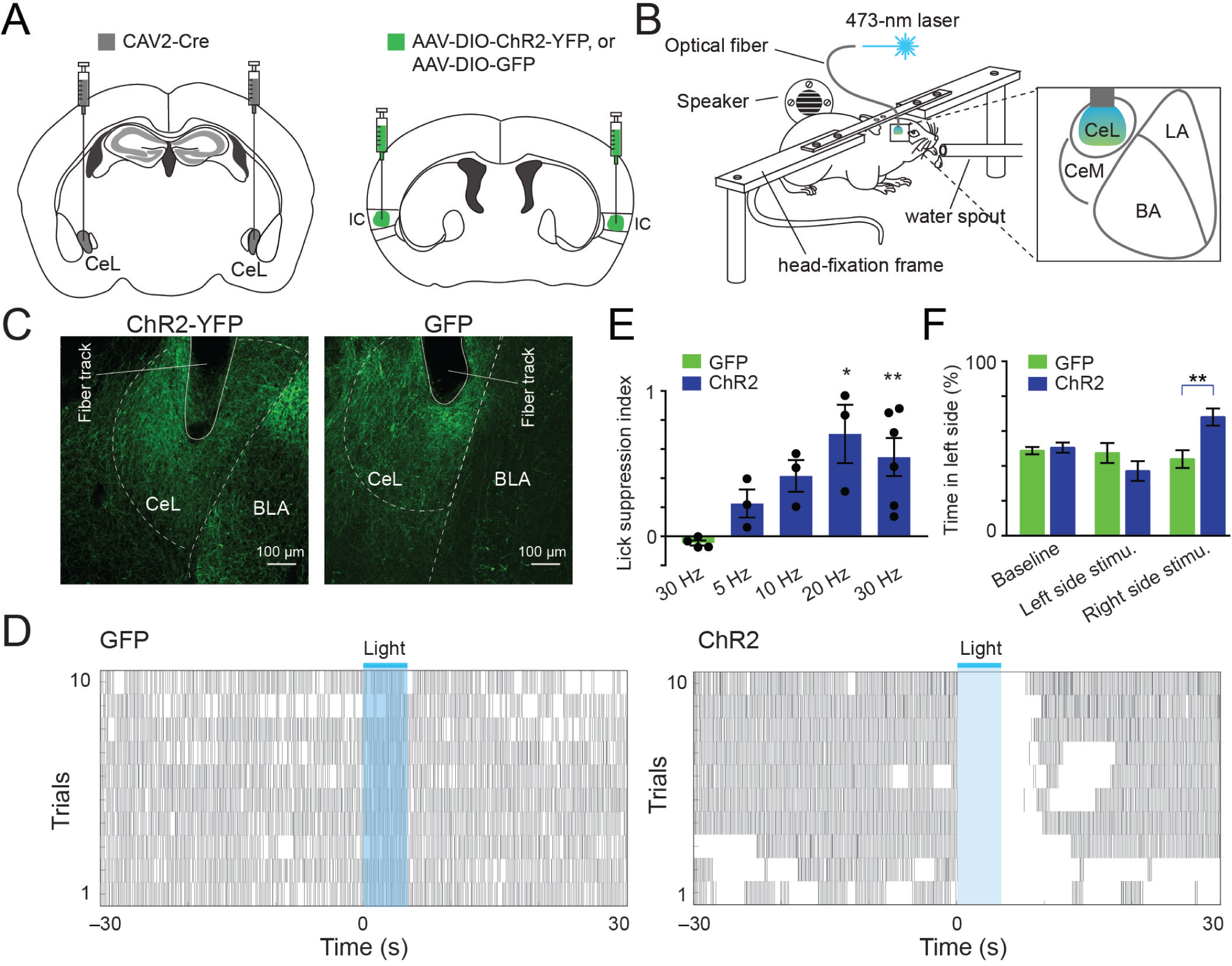
Optogenetic activation of the IC-CeL circuit is sufficient to induce action suppression and avoidance behavior. (**A** & **B**) Schematics of the experimental design. (**C**) Representative images showing the locations of optical fiber implantation and green axon fibers in the CeL, which originated from CeL-projecting IC neurons expressing ChR2-YFP (left) or GFP (right). (**D**) Raster plots showing licking behavior for a GFP (left) and a ChR2 (right) animal. Blue bars and shaded areas indicate the time window of laser stimulation. (**E**) Quantification of the effect of the photostimulation on licking behavior (suppression index) at different stimulation frequencies (F(4,14) = 4.375, *P* = 0.017; \**P* = 0.024, \*\**P* = 0.007; one-way ANOVA followed by Sidak’s multiple comparisons test; 5-20Hz, n = 3 mice; 30Hz, n = 6 mice). (**F**) In a real-time place avoidance behavioral paradigm, the ChR2 animals avoided the stimulation side (two-way RM ANOVA main effect of stimulation side, F(2,16) = 4.27; *P* = 0.03; main effect of virus treatment, F(1,8) = 1.53; *P* = 0.25); interaction, F(2,16) = 6.94; *P* = 0.007; \*\**P* = 0.005, Post hoc Sidak’s multiple comparisons test; GFP, n = 4 mice, ChR2, n = 6 mice).

Four to five weeks following surgery, these mice were water deprived and trained to achieve stable licking to a water spout, during which we delivered pulses of blue light into the CeL. Photostimulation in the CeL in which the axon terminals originating from the IC expressed ChR2 elicited robust suppression of licking, an effect that was dependent on the frequency of stimulation (Fig. 5D, E). Furthermore, such optogenetic activation of the IC-CeL pathway induced place aversion in a real time place aversion (RTPA) task (Fig. 5F). By contrast, photostimulation in the CeL in which IC axons expressed GFP induced neither lick suppression nor place aversion (Fig. 5D-F). These results suggest that activation of the IC-CeL circuit is aversive, and is sufficient to induce action suppression and avoidance responses.

To test whether activation of the IC-CeL circuit can be substituted for an aversive tastant to instruct avoidance learning, similar to learning of the no-go response in the go/no-go task, we trained the same mice as those used in Fig. 5 in a modified go/no-go task, in which licking during one cue led to water delivery alone (the “laser-off” trials), whereas licking during another cue resulted in water delivery coinciding with bilateral photostimulation in the CeL (the “laser-on” trials) (Fig. 6A; also see Methods). We found that the mice in which the IC-CeL pathway expressed ChR2 – and thus could be activated by the photostimulation – showed reduced responding to the cue in the laser-on trials compared with in the laser-off trials (Fig. 6B, C). By contrast, the mice in which the IC-CeL pathway expressed GFP showed similar cue-evoked responding in the laser-on or laser-off trials (Fig. 6B, C). We also verified that the photostimulation did not cause obvious motor effects in either the ChR2 mice or the GFP mice in an open field setting (Fig. 6D).

**Figure 6.**
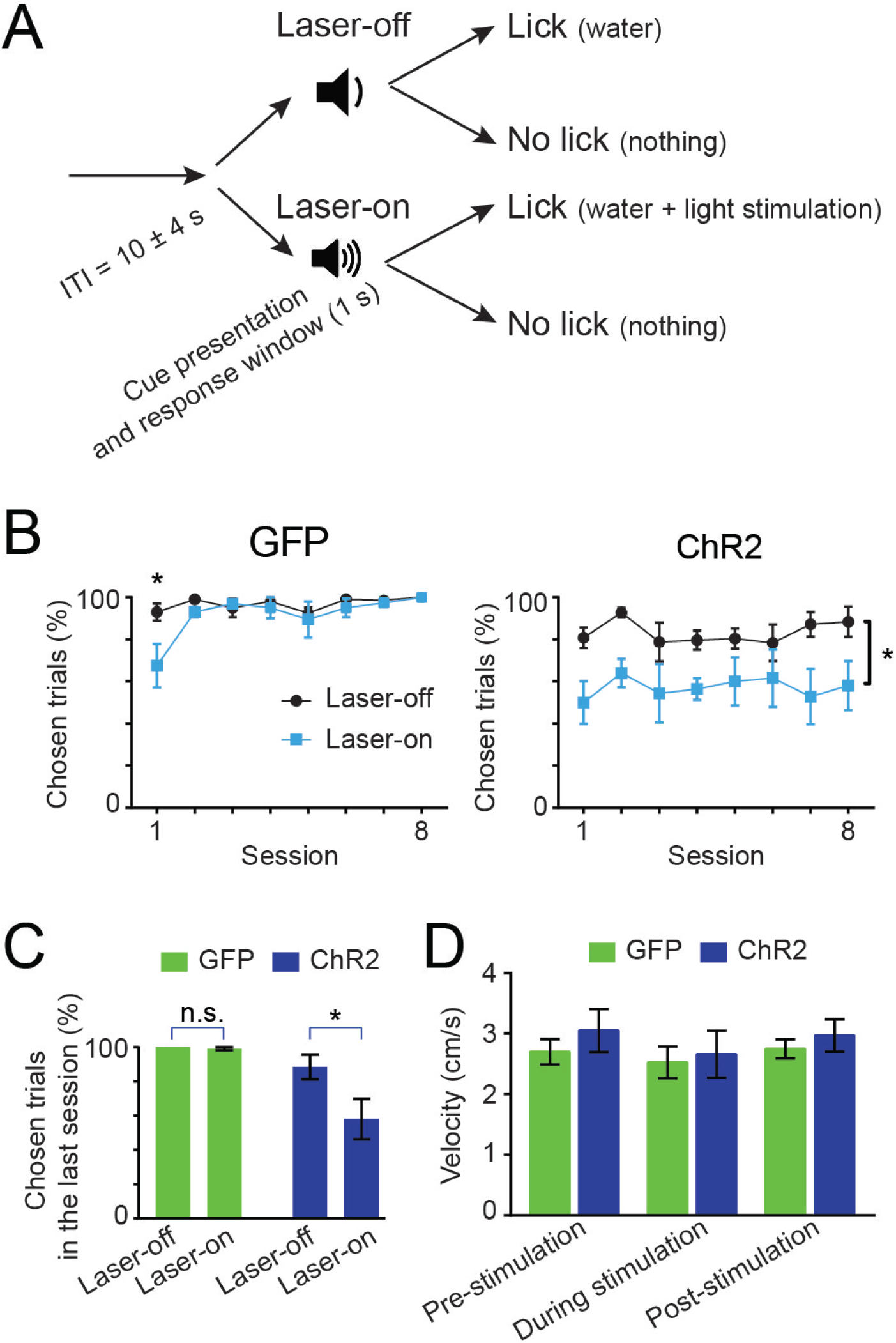
Optogenetic activation of the IC-CeL circuit is sufficient to instruct avoidance learning. (**A**) A schematic of the experimental design. (**B**) Animal behavior is quantified as the percentage of trials that animals chose to lick during CS presentation (chosen trials) (GFP animals: two-way RM ANOVA showed a main effect of session (F(5,30) = 5.59; *P* < 0.001, no main effect of trial type (F(1,6) = 2.07; *P* = 0.2, and a significant interaction (F(5,30) = 3.05; *P* = 0.024; Post hoc Sidak’s multiple comparisons tests revealed a significant difference in session 1 (*P* = 0.004, n = 4 mice). ChR2 mice: two-way RM ANOVA showed a main effect of trial type (F(1,8) = 5.7; *P* = 0.04), no main effect of session (F(7,56) = 1.35; *P* = 0.25) or interaction (F(7,56) = 0.71; *P* = 0.66; n = 5 mice). (**C**) Quantification of the chosen trials in the final training session (two-way repeated measures ANOVA revealed a main effect of group (GFP vs. ChR2) (F(1,7) = 8.19; *P* = 0.024), a main effect of trial type (laser-off vs. laser-on) (F(1,7) = 6.59; *P* = 0.037), and a significant interaction (F(1,7) = 5.778; *P* = 0.047); \**P* = 0.015, Post hoc Sidak’s multiple comparisons test). (**D**) In an open field, laser stimulation did not affect movement of mice (GFP or ChR2 mice), measured as average movement velocity over 10 trials (*P* > 0.05, two-way RM ANOVA). All data are presented as mean ± s.e.m.

Of note, although optogenetic activation of the IC-CeL pathway was less effective than quinine reinforcement, we observed a pattern of responding in these mice qualitatively similar to that of the control animals trained on the standard sucrose/quinine go/no-go task (Fig. 4B), in which the mice showed low levels of responding to the no-go cue and high levels of responding to the go cue. Together, these results indicate that activation of the IC-CeL pathway is sufficient to instruct the learning of anticipatory action suppression. This mechanism is likely engaged when learning to avoid an aversive outcome, such as an unpleasant tastant.

## Discussion

In this study we examined the role of the IC-CeL circuit in the establishment of behavioral inhibition for avoiding an unpleasant tastant. Using retrograde anatomic tracing approaches, including the modified rabies virus-assisted tracing, together with ChR2-based circuit mapping, we found that the GC region of the IC sends direct excitatory projections to the CeL, which make monosynaptic connections with both the SOM^+^ and SOM^−^ CeL neurons, with the latter being mainly PKC-δ^+^ neurons (Li et al., 2013; Penzo et al., 2014). Specific inhibition of the CeL-projecting IC neurons with TeLC prevented mice from acquiring the no-go response, and only mildly affected the go response in a tastant (sucrose/quinine)-reinforced go/no-go task. Furthermore, selective activation of the IC-CeL pathway with optogenetics drove unconditioned lick suppression in thirsty animals, induced avoidance behavior, and was sufficient to instruct conditioned action suppression in response to the cue that predicts the optogenetic activation. These results demonstrate that activity in the IC-CeL circuit is necessary for establishing anticipatory avoidance responses to an unpleasant tastant, and is also sufficient to drive learning of such anticipatory avoidance responses.

The IC has been shown to contain regions with selective responses to distinct tastants (Chen et al., 2011). As measured in anesthetized animals, these “hotspots” occupy non-overlapping regions within the GC. The region with preferential responses to bitter tastes resides in the posterior part of the GC, which corresponds approximately to the area of IC we identified that sends the strongest projections to the CeL (also see (Allen et al., 1991)). These findings suggest that the IC may preferentially convey information about bitter/aversive tastants to the CeL, which in turn drive behavioral inhibition. Interestingly, although the IC has been shown to be required for identifying and discriminating between tastants (Katz et al., 2001; Peng et al., 2015; Sammons et al., 2016), we did not observe an effect on quinine sensitivity when we selectively silenced the IC-CeL pathway with TeLC (Fig. 4F), suggesting that the ability for bitter tastant identification and that for processing the basic reinforcing properties of bitter tastants are preserved in these mice, at least when cognitive demand is low.

The CeA, including the CeL, receives aversive information of different modalities directly from the brainstem parabrachial nucleus (PBN) (Carter et al., 2013; Han et al., 2015; Norgren et al., 1989; Sato et al., 2015). Previous studies have shown that the CeA responds to noxious stimuli, such as footshocks that induce somatic pain (Han et al., 2015; Radulovic et al., 1998), colorectal distension that induces visceral pain (Myers and Greenwood-Van Meerveld, 2012), and lithium chloride, which induces malaise and is the most commonly used US for inducing conditioned taste aversion (Lamprecht and Dudai, 1995). The CeA has also been implicated in taste processing (Sadacca et al., 2012) and feeding behaviors (Cai et al., 2014). Thus, the CeA is anatomically poised to process convergent somatosensory, visceral, gustatory and aversive information and may be recruited by multiple neural circuits for action suppression in a variety of tasks, in which the goal is to avoid unwanted consequences.

More recent studies, including those of our own, indicate that the CeA contains functionally heterogeneous neuronal populations. SOM^+^ CeL neurons control passive defensive responses, such as freezing and action suppression (Fadok et al., 2017; Li et al., 2013; Penzo et al., 2014; Penzo et al., 2015; Yu et al., 2016), whereas PKC-δ^+^ CeL neurons are involved in conveying aversive US information and instructing aversive learning (Yu et al., 2017). The PKC-δ^+^ CeL neurons have also been linked to suppression of feeding (Cai et al., 2014). As both of these CeL populations receive direct excitatory inputs from the IC (Fig. 1), they may contribute to distinct aspects of the IC-CeL circuit function described in the current study. An intriguing possibility is that the SOM^+^ population induces action suppression when excited by IC inputs, while the PKC-δ^+^ population regulates learning and potentially aversion when activated by the same inputs. Some of the functions mediated by CeL neurons, in particular those by PKC-δ^+^ neurons, are consistent with the findings that the CeA plays a role in alerting or attentional processes, and can explain our observations suggesting that the IC-CeL circuit may have a more general function in behavior, i.e., it influences performance and actions during not only the no-go trials, but also the go trials of the go/no-go task, although the impact on the no-go trials is much stronger.

## Acknowledgements

We thank A. Fontanini (Stony Brook University, USA) for critical reading of an early version of the manuscript, J. Johansen (RIKEN Brain Science Institute, Japan) for helpful discussions, P. Wulff (University of Kiel, Germany) for kindly providing the AAV.CAG.Flex.TeLC-eGFP plasmid, and members of the Li laboratory for discussions. This work was supported by grants from NARSAD (23169, B.L.), National Natural Science Foundation of China (81428010, B.L. and M.H.), National Institutes of Mental Health (R01MH101214, B.L.), Wodecroft Foundation (B.L.), Stanley Family Foundation (B.L.), Simons Foundation Autism Research Initiative (344904, SFARI) (B.L.), and Human Frontier Science Program (RGP0015, B.L.).

## References

Accolla, R., and Carleton, A. (2008). Internal body state influences topographical plasticity of sensory representations in the rat gustatory cortex. Proc Natl Acad Sci U S A 105, 4010–4015.

Ahrens, S., Jaramillo, S., Yu, K., Ghosh, S., Hwang, G.R., Paik, R., Lai, C., He, M., Huang, Z.J., and Li, B. (2015). ErbB4 regulation of a thalamic reticular nucleus circuit for sensory selection. Nat Neurosci 18, 104–111.

Allen, G.V., Saper, C.B., Hurley, K.M., and Cechetto, D.F. (1991). Organization of visceral and limbic connections in the insular cortex of the rat. J Comp Neurol 311, 1–16.

Balleine, B.W. (2005). Neural bases of food-seeking: affect, arousal and reward in corticostriatolimbic circuits. Physiol Behav 86, 717–730.

Balleine, B.W. (2011). Sensation, Incentive Learning, and the Motivational Control of Goal-Directed Action. In Neurobiology of Sensation and Reward, J.A. Gottfried, ed. (Boca Raton (FL)).

Balleine, B.W., and Killcross, S. (2006). Parallel incentive processing: an integrated view of amygdala function. Trends in Neurosciences 29, 272–279.

Bermudez-Rattoni, F. (2004). Molecular mechanisms of taste-recognition memory. Nat Rev Neurosci 5, 209–217.

Bru, T., Salinas, S., and Kremer, E.J. (2010). An update on canine adenovirus type 2 and its vectors. Viruses 2, 2134–2153.

Cai, H., Haubensak, W., Anthony, T.E., and Anderson, D.J. (2014). Central amygdala PKC-delta(+) neurons mediate the influence of multiple anorexigenic signals. Nat Neurosci 17, 1240–1248.

Callaway, E.M., and Luo, L. (2015). Monosynaptic Circuit Tracing with Glycoprotein-Deleted Rabies Viruses. J Neurosci 35, 8979–8985.

Carter, M.E., Soden, M.E., Zweifel, L.S., and Palmiter, R.D. (2013). Genetic identification of a neural circuit that suppresses appetite. Nature 503, 111–114.

Chen, X., Gabitto, M., Peng, Y., Ryba, N.J., and Zuker, C. S. (2011). A gustotopic map of taste qualities in the mammalian brain. Science 333, 1262–1266.

Ciocchi, S., Herry, C., Grenier, F., Wolff, S.B., Letzkus, J.J., Vlachos, I., Ehrlich, I., Sprengel, R., Deisseroth, K., Stadler, M.B., et al. (2010). Encoding of conditioned fear in central amygdala inhibitory circuits. Nature 468, 277–282.

Fadok, J.P., Krabbe, S., Markovic, M., Courtin, J., Xu, C., Massi, L., Botta, P., Bylund, K., Muller, C., Kovacevic, A., et al. (2017). A competitive inhibitory circuit for selection of active and passive fear responses. Nature 542, 96–100.

Gallagher, M., Graham, P.W., and Holland, P.C. (1990). The amygdala central nucleus and appetitive Pavlovian conditioning: lesions impair one class of conditioned behavior. J Neurosci 10, 1906–1911.

Gardner, M.P., and Fontanini, A. (2014). Encoding and tracking of outcome-specific expectancy in the gustatory cortex of alert rats. J Neurosci 34, 13000–13017.

Goosens, K.A., and Maren, S. (2003). Pretraining NMDA receptor blockade in the basolateral complex, but not the central nucleus, of the amygdala prevents savings of conditional fear. Behav Neurosci 117, 738–750.

Han, S., Soleiman, M.T., Soden, M.E., Zweifel, L.S., and Palmiter, R.D. (2015). Elucidating an Affective Pain Circuit that Creates a Threat Memory. Cell 162, 363–374.

Haney, R.Z., Calu, D.J., Takahashi, Y.K., Hughes, B.W., and Schoenbaum, G. (2010). Inactivation of the central but not the basolateral nucleus of the amygdala disrupts learning in response to overexpectation of reward. J Neurosci 30, 2911–2917.

Hashimoto, K., and Spector, A.C. (2014). Extensive lesions in the gustatory cortex in the rat do not disrupt the retention of a presurgically conditioned taste aversion and do not impair unconditioned concentration-dependent licking of sucrose and quinine. Chem Senses 39, 57–71.

Haubensak, W., Kunwar, P.S., Cai, H., Ciocchi, S., Wall, N.R., Ponnusamy, R., Biag, J., Dong, H.-W., Deisseroth, K., Callaway, E.M., et al. (2010). Genetic dissection of an amygdala microcircuit that gates conditioned fear. Nature 468, 270–276.

Katz, D.B., Simon, S.A., and Nicolelis, M.A. (2001). Dynamic and multimodal responses of gustatory cortical neurons in awake rats. J Neurosci 21, 4478–4489.

Kelley, A.E. (2004). Ventral striatal control of appetitive motivation: role in ingestive behavior and reward-related learning. Neurosci Biobehav Rev 27, 765–776.

Kentridge, R.W., Shaw, C., and Aggleton, J.P. (1991). Amygdaloid lesions and stimulus-reward associations in the rat. Behav Brain Res 42, 57–66.

Kim, J., Zhang, X., Muralidhar, S., LeBlanc, S.A., and Tonegawa, S. (2017). Basolateral to Central Amygdala Neural Circuits for Appetitive Behaviors. Neuron 93, 1464–1479 e1465.

Kusumoto-Yoshida, I., Liu, H., Chen, B.T., Fontanini, A., and Bonci, A. (2015). Central role for the insular cortex in mediating conditioned responses to anticipatory cues. Proc Natl Acad Sci U S A 112, 1190–1195.

Lamprecht, R., and Dudai, Y. (1995). Differential modulation of brain immediate early genes by intraperitoneal LiCl. Neuroreport 7, 289–293.

Li, H., Penzo, M.A., Taniguchi, H., Kopec, C.D., Huang, Z.J., and Li, B. (2013). Experience-dependent modification of a central amygdala fear circuit. Nature Neuroscience 16, 332–339.

Li, L., Tasic, B., Micheva, K.D., Ivanov, V.M., Spletter, M.L., Smith, S.J., and Luo, L. (2010). Visualizing the distribution of synapses from individual neurons in the mouse brain. PLoS One 5, e11503.

Madisen, L., Zwingman, T.A., Sunkin, S.M., Oh, S.W., Zariwala, H.A., Gu, H., Ng, L.L., Palmiter, R.D., Hawrylycz, M.J., Jones, A.R., et al. (2010). A robust and high-throughput Cre reporting and characterization system for the whole mouse brain. Nature Neuroscience 13, 133–140.

Murray, A.J., Sauer, J.F., Riedel, G., McClure, C., Ansel, L., Cheyne, L., Bartos, M., Wisden, W., and Wulff, P. (2011). Parvalbumin-positive CA1 interneurons are required for spatial working but not for reference memory. Nat Neurosci 14, 297–299.

Myers, B., and Greenwood-Van Meerveld, B. (2012). Differential involvement of amygdala corticosteroid receptors in visceral hyperalgesia following acute or repeated stress. Am J Physiol Gastrointest Liver Physiol 302, G260–266.

Norgren, R., Nishijo, H., and Travers, S.P. (1989). Taste responses from the entire gustatory apparatus. Ann N Y Acad Sci 575, 246–263; discussion 263-244.

Peng, Y., Gillis-Smith, S., Jin, H., Trankner, D., Ryba, N.J., and Zuker, C.S. (2015). Sweet and bitter taste in the brain of awake behaving animals. Nature 527, 512–515.

Penzo, M.A., Robert, V., and Li, B. (2014). Fear conditioning potentiates synaptic transmission onto long-range projection neurons in the lateral subdivision of central amygdala. The Journal of neuroscience: the official journal of the Society for Neuroscience 34, 2432–2437.

Penzo, M.A., Robert, V., Tucciarone, J., De Bundel, D., Wang, M., Van Aelst, L., Darvas, M., Parada, L.F., Palmiter, R.D., He, M., et al. (2015). The paraventricular thalamus controls a central amygdala fear circuit. Nature 519, 455–459.

Petrovich, G.D., Ross, C.A., Mody, P., Holland, P.C., and Gallagher, M. (2009). Central, but not basolateral, amygdala is critical for control of feeding by aversive learned cues. J Neurosci 29, 15205–15212.

Radulovic, J., Kammermeier, J., and Spiess, J. (1998). Relationship between fos production and classical fear conditioning: effects of novelty, latent inhibition, and unconditioned stimulus preexposure. The Journal of neuroscience: the official journal of the Society for Neuroscience 18, 7452–7461.

Robinson, M.J., Warlow, S.M., and Berridge, K.C. (2014). Optogenetic excitation of central amygdala amplifies and narrows incentive motivation to pursue one reward above another. J Neurosci 34, 16567–16580.

Roesch, M.R., Esber, G.R., Li, J., Daw, N.D., and Schoenbaum, G. (2012). Surprise! Neural correlates of Pearce-Hall and Rescorla-Wagner coexist within the brain. Eur J Neurosci 35, 1190–1200.

Sadacca, B.F., Rothwax, J.T., and Katz, D.B. (2012). Sodium concentration coding gives way to evaluative coding in cortex and amygdala. J Neurosci 32, 9999–10011.

Sammons, J., Bass, C., Victor, J., and Di Lorenzo, P. (2016). Gustatory cortical input onto the nucleus of the solitary tract refines neuronal firing patterns and enhances learning in the awake rat. Society for Neuroscience; 2016; San Diego, CA, USA.

Samuelsen, C.L., and Fontanini, A. (2017). Processing of Intraoral Olfactory and Gustatory Signals in the Gustatory Cortex of Awake Rats. J Neurosci 37, 244–257.

Sato, M., Ito, M., Nagase, M., Sugimura, Y.K., Takahashi, Y., Watabe, A.M., and Kato, F. (2015). The lateral parabrachial nucleus is actively involved in the acquisition of fear memory in mice. Mol Brain 8, 22.

Seo, D.O., Funderburk, S.C., Bhatti, D.L., Motard, L.E., Newbold, D., Girven, K.S., McCall, J.G., Krashes, M., Sparta, D.R., and Bruchas, M.R. (2016). A GABAergic Projection from the Centromedial Nuclei of the Amygdala to Ventromedial Prefrontal Cortex Modulates Reward Behavior. J Neurosci 36, 10831–10842.

Stephenson-Jones, M., Yu, K., Ahrens, S., Tucciarone, J.M., van Huijstee, A.N., Mejia, L.A., Penzo, M.A., Tai, L.H., Wilbrecht, L., and Li, B. (2016). A basal ganglia circuit for evaluating action outcomes. Nature 539, 289–293.

Taniguchi, H., He, M., Wu, P., Kim, S., Paik, R., Sugino, K., Kvitsiani, D., Fu, Y., Lu, J., Lin, Y., et al. (2011). A resource of Cre driver lines for genetic targeting of GABAergic neurons in cerebral cortex. Neuron 71, 995–1013.

Vincis, R., and Fontanini, A. (2016). Associative learning changes cross-modal representations in the gustatory cortex. Elife 5.

Wilensky, A.E., Schafe, G.E., Kristensen, M.P., and LeDoux, J.E. (2006). Rethinking the fear circuit: the central nucleus of the amygdala is required for the acquisition, consolidation, and expression of Pavlovian fear conditioning. The Journal of neuroscience: the official journal of the Society for Neuroscience 26, 12387–12396.

Yamamoto, T., Yuyama, N., Kato, T., and Kawamura, Y. (1985). Gustatory responses of cortical neurons in rats. II. Information processing of taste quality. J Neurophysiol 53, 1356–1369.

Yasoshima, Y., and Yamamoto, T. (1998). Short-term and long-term excitability changes of the insular cortical neurons after the acquisition of taste aversion learning in behaving rats. Neuroscience 84, 1–5.

Yu, K., Ahrens, S., Zhang, X., Schiff, H., Ramakrishnan, C., Fenno, L., Deisseroth, K., Zhou, P., Paninski, L., and Li, B. (2017). The central amygdala controls learning in the lateral amygdala. bioRxiv 126649; doi: https://doiorg/101101/126649.

Yu, K., Garcia da Silva, P., Albeanu, D.F., and Li, B. (2016). Central Amygdala Somatostatin Neurons Gate Passive and Active Defensive Behaviors. J Neurosci 36, 6488–6496.

Zhang, F., Wang, L.P., Boyden, E.S., and Deisseroth, K. (2006). Channelrhodopsin-2 and optical control of excitable cells. Nat Methods 3, 785–792.

